# Finding and Following: A deep learning-based pipeline for tracking platelets during thrombus formation *in vivo* and *ex vivo*

**DOI:** 10.1101/2023.10.02.560609

**Authors:** Abigail S. McGovern, Pia Larsson, Volga Tarlac, Natasha Setiabakti, Leila Shabani Mashcool, Justin R. Hamilton, Niklas Boknäs, Juan Nunez-Iglesias

**Affiliations:** Biomedicine Discovery Institute, Monash University, Melbourne, Australia; Australian Centre for Blood Diseases, Monash University, Melbourne, Australia; Department of Biomedical and Clinical Sciences Linköping University, Linköping, Sweden

**Keywords:** thrombosis, platelets, segmentation, deep learning, tracking, image analysis

## Abstract

The last decade has seen increasing use of advanced imaging techniques in platelet research. However, there has been a lag in the development of image analysis methods, leaving much of the information trapped in images. Herein, we present a robust analytical pipeline for finding and following individual platelets over time in growing thrombi. Our pipeline covers four steps: detection, tracking, estimation of tracking accuracy, and quantification of platelet metrics. We detect platelets using a deep learning network for image segmentation, which we validated with proofreading by multiple experts. We then track platelets using a standard particle tracking algorithm and validate the tracks with custom image sampling — essential when following platelets within a dense thrombus. We show that our pipeline is more accurate than previously described methods. To demonstrate the utility of our analytical platform, we use it to show that *in vivo* thrombus formation is much faster than that *ex vivo*. Furthermore, platelets *in vivo* exhibit less passive movement in the direction of blood flow. Our tools are free and open source and written in the popular and user-friendly Python programming language. They empower researchers to accurately find and follow platelets in fluorescence microscopy experiments.x

**Plain language summary:** In this paper we describe computational tools to find and follow individual platelets in blood clots recorded with fluorescence microscopy. Our tools work in a diverse range of conditions, both in living animals and in artificial flow chamber models of thrombosis. Our work uses deep learning methods to achieve excellent accuracy. We also provide tools for visualising data and estimating error rates, so you don’t have to just trust the output. Our workflow measures platelet density, shape, and speed, which we use to demonstrate differences in the kinetics of clotting in living vessels versus a synthetic environment. The tools we wrote are open source, written in the popular Python programming language, and freely available to all. We hope they will be of use to other platelet researchers.

## Introduction

The last decade has seen an explosion in the usage of modern imaging modalities to capture the behaviour of platelets in growing blood clots. Using fluorescence microscopy, platelet biologists can now capture time resolved three-dimensional data both *in vivo* and *ex vivo*. However, quantitative methods lag these advances in imaging, leaving important biological insights trapped in image data. In the ideal case, we aim to collect data on each individual platelet over time. Early attempts to do this used labour-intensive manual platelet identification and tracking [1,2]. Although they provided interesting insights into platelet behaviour, such manual methods allowed only a small number of platelets to be followed for short periods of time. Understanding platelet behaviour in the context of the whole thrombus requires large scale automatic platelet detection and tracking.

One of the earliest uses of such automated detection in platelet research was by Claesson, Lindahl and Faxälv [3], who used the classical Difference of Gaussian (DoG) blob detection algorithm to detect and count platelets in images of *ex vivo* thrombosis. They showed that counting platelets was a more robust measure of thrombus dynamics than measuring total fluorescence volume or intensity. However, classical algorithms are not robust to spatial variations in fluorescence intensity, and require careful parameter tuning when imaging conditions change [4]. Indeed, in our preliminary live animal thrombosis experiments, we found classical segmentation algorithms to be brittle: what worked for one condition failed for another. We hypothesized that a machine learning approach would be more robust.

Deep learning methods can segment objects with near human precision [5]. Although deep leaning-based segmentation has been applied to identifying platelets in 2D histology imaging [6,7], it has never been used to identify platelets in 3D. A deep network can learn sophisticated rules to detect objects even in noisy data. Once cells are detected in every frame, we can use a tracking algorithm [8] to follow them over time.

In this study, we present a deep learning-based pipeline to analyse 3D timelapse fluorescence microscopy images of platelets in growing thrombi. Our pipeline identifies and tracks large numbers of individual platelets during thrombosis, whether *in vivo* or *ex vivo*. We use a U-net-based [9] segmentation algorithm (*PlateSeg*), followed by a classical approach for tracking objects. As the size of these datasets makes it challenging to validate or even visualise our results [10], we also developed methods and tools for rapid assessment of tracking accuracy.

Our analysis provides time-resolved quantifications of the fluorescence intensity, movement, shape, and size of each platelet identified from image data. Our workflow is more accurate and robust to noise than previously used methods and is able to segment platelets with different types of labels. To demonstrate the utility of this method, we report fundamental differences in the speed and directionality of platelet movement between *in vivo* and *ex vivo* experimental models of thrombus formation. We have made our quality-assurance and quantitative methods available through several graphical user interface (GUI)-based tools that we continue to improve.

## Methods

### In vivo thrombosis experiments

All animal procedures were approved by the Alfred Medical Research and Education Precinct Animal Ethics Committee (AMREP Ethics approval number E/1912/2019/M). Our scanning laser ablation mesenteric venule thrombosis model (scanning-LIEI) has been previously described in detail [11]. To obtain fractionally labelled platelets, mice used for intravital imaging were injected with 30 × 10^6^ labelled donor platelets resulting in approximately 2.5% labelled platelets circulating in the mouse. Donor platelets were prepared as described in Maxwell, Yuan [12] and labelled with Dylight-649-anti-GPIbB (Emfret Analytics, product # X-649).

### Ex vivo thrombosis experiments

Mouse blood was collected in lepirudin (final concentration 80 U/ml) from the inferior vena cava of terminally anaesthetized (Sodium Pentobarbitone 90 mg/kg) mice and used for experimentation within 30 min of collection. A fraction of blood, corresponding to 2.5% of the total volume, was removed, labelled with platelet marker X-649 for 10 min and mixed back into the blood to obtain a fraction of labelled platelets of approximately 2.5%.

Human blood collection was approved by the Monash University Human Research Ethics committee (project number 2017-9712-13995). After informed consent, blood from healthy volunteers was collected by venepuncture using enoxaparin as anticoagulant (final concentration 40 U/ml) and used for experimentation within 2h of collection. Fractional labelling of platelets was obtained as described for mouse blood but labelled with an APC-labelled anti-human CD41 antibody (Abcam, product # ab55866).

A polydimethylsiloxane (PDMS) microfluidic device consisting of straight channels of 1000 μm width and 100 μm height was placed on a #1 cover slip and the glass surface coated with type I collagen (Chronolog, 200 μg/ml) for 30 min and subsequently blocked with BSA (2 mg/ml) for 20 min prior to mouse or human blood perfusion driven by a syringe pump (Harvard Apparatus). Blood was perfused at wall shear rates of 600 s^-1^, 1800 s^-1^ or 3000 s^-1^ for up to 10 min.

### Imaging

Our Nikon A1R+ imaging platform has been described in detail [11] and was utilized both for *in vivo* and *ex vivo* imaging. Images collected were of 512 × 512 pixels, scan speed of 30 fps, scanner zoom of 2x (250 μm × 250 μm), bidirectional scanning, and 2 x line averaging. Emitted fluorescence was detected with a pinhole size of 57.5 μm. The piezo Z-stage was used to capture continuous Z-stack images with a Z-axis step size of 2 μm and a Z-range of 64 μm (time to acquire one Z-stack 3.1s) for up to10 min.

### Deep learning segmentation

A modified U-net architecture, as originally described in [29], is used as the backbone for deep-learning based instance segmentation. The network was trained with the Adam optimiser [13] at a learning rate of 0.01. Training data consisted of randomly generated sub-volumes extracted from ground truth segmentations. From each of 10 ground truth frames, we extracted 50 volumes, with dimensions of 10 × 256 × 256 voxels (500 batches, mini batch size = 1). Prior to training, data was augmented with a number of geometric and intensity transformations. The network was trained to produce feature maps that could be processed to produce object labels. The feature maps included object borders in each axis (1^st^ order edge affinities for x, y, z axes), centre point prediction, and a mask (image foreground). To predict object centres, we computed ‘centredness’ which is the normalised (by maximum value) inverse distance from the centre point. Feature maps are fed to custom modified watershed algorithm to generate platelet labels. The network and algorithm was implemented using pytorch [14] and is available via the napari [15] GUI plugin Iterseg (https://github.com/AbigailMcGovern/iterseg, DOI: 10.5281/zenodo.10572073), which is under continued development to facilitate others to train their own network. Installation for this plugin can be seen in Figure 1.A and an image of the user interface can be seen in Figure 1.B.

**Figure 1.**
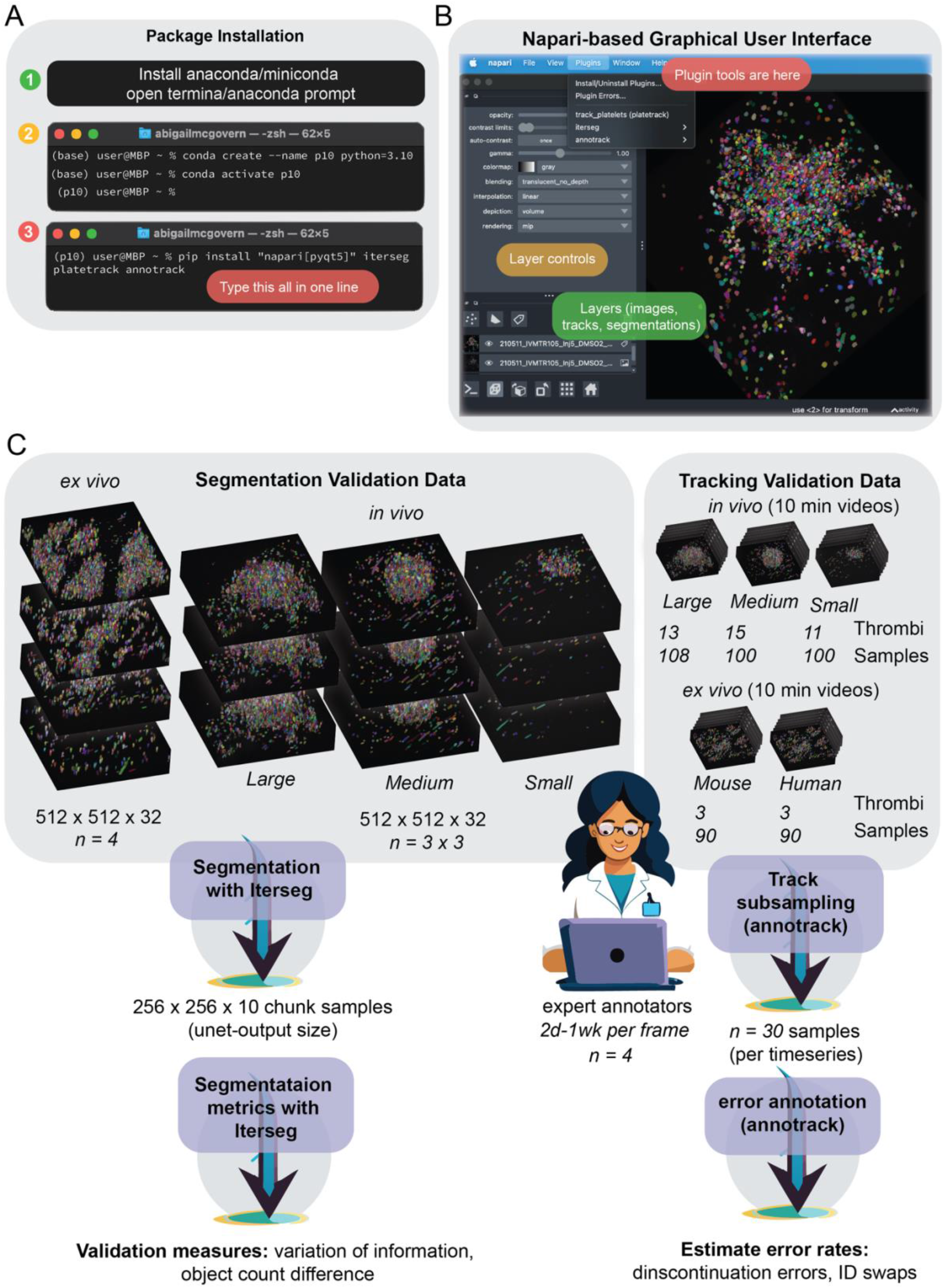
Overview of software installation, interface, and methods validation. (**A**) Our software can be installed in three steps. Whilst it is possible to install our software using only python and pip (step 3), it is ideal to install anaconda or miniconda (step 1) and generate a new environment (step 2). Once in the new environment, packages can be installed in only one line (step 3) and opened by typing “napari” into the command line. (**B**) An image showing the napari user interface and how to access our software through the plugins menu. (**C**) Schematic outlining the data and the process used to validate our methodology.

### Ground truth data

Ground truth data was produced by correcting a segmentation produced by an earlier version of the neural network using the napari [15] plugin Zarpaint (source code at https://github.com/jni/zarpaint), which we developed alongside this project. For training and validation data we had four people proof-read data to produce ground truth.

### Segmentation validation

Validation of segmentation accuracy involved assessing the quality of segmentation in unseen volumes (volumes not used in tracking) for which ground truth data was created. We used unseen images of three thrombi of different sizes to validate the segmentation and had three expert scientists generate ground truth for each thrombus (Figure 1.C), meaning that each person proofread ∼4000 platelets. The small thrombus was treated with cangrelor to represent a treatment condition. The medium and large thrombi were generated in mice injected with saline or 20% DMSO, respectively. Assessments included variation of information (VI) and the difference in platelet count. VI is the sum of two asymmetrical sub-scores denoting the conditional entropy between segmentations. These sub-scores represent under- and over-segmentation. VI was assessed using the Python module Scikit Image [16]. Segmentation accuracy assessment is available via the Napari plugin Iterseg.

### Data tabulation and tracking

Prior to tracking, platelet data was extracted from segmentations and tabulated using the Python package scikit-image [16]. We extracted data about platelet elongation and flatness (as described in Kong and Fonseca [17]), fluorescence intensity, and volume. Tracking was performed using the Python package TrackPy [18]. TrackPy uses an algorithm pioneered by Cocker and Grier [8], which assumes objects move with Brownian motion and employs a tracking-by-detection paradigm to find a globally optimal solution for every time step. Following tracking, we estimate velocity in each axis using finite difference derivatives. This method has been made available through a user interface as a napari plugin (https://github.com/AbigailMcGovern/platelet-tracking, DOI: 10.5281/zenodo.10572093). Installation for this plugin can be seen in Figure 1.A.

### Tracking validation

We estimated tracking accuracy by sampling tracks, which are then assessed for errors by a human proofreader. We randomly sampled subsections of platelet tracks (∼31 frames). Track segments were annotated for errors using annotrack, a napari plugin we developed (https://github.com/AbigailMcGovern/annotrack, DOI: 10.5281/zenodo.10572091). For each track segment, both identity swaps and track discontinuations were annotated. Error rates were defined according to the number of errors per frame.

### Statistical and graphical methods

Statistics were done using the Python library scipy’s [19] stats module and comprised 95% confidence intervals, Kruskal-Wallis tests (reported as H-values and p-values), and Mann Whitney U-tests (reported as U-values and p-values). Plots used the Python libraries seaborn, matplotlib [20], and ptitprince [21]. Paraview software [22] was used to render platelet coordinates for visualisations.

### Software and data availability

Our software is available to install from PyPI as the packages iterseg, platetrack, and annotrack. Archived versions matching the versions described in this paper are available as Zenodo IDs DOI: 10.5281/zenodo.10633360, DOI: 10.5281/zenodo.10572093, and DOI: 10.5281/zenodo.10572091. Source code is available at GitHub repos https://github.com/AbigailMcGovern/iterseg, https://github.com/AbigailMcGovern/platelet-tracking, and https://github.com/AbigailMcGovern/annotrack, and contributions from interested readers are welcome. Example data is available on Figshare at DOI: 10.26180/25137497. Instructions for software use can be found in the Supplementary material.

## Results

### Finding platelets in vivo and ex vivo with deep learning

To assess the accuracy of our automated platelet finding, we compared our results with manual platelet segmentations performed by three trained biologists *(“ground truth”*). The validation process is outlined in Figure 1.A. For our *in vivo* data, we randomly selected frames from thrombi of different sizes. To provide context, we compared our trained network, *PlateSeg*, with the previously used DoG approach. **Figure 2.A** shows the human-generated labels, PlateSeg labels, and DoG labels for a representative thrombus. PlateSeg makes few mistakes with regards to the ground truth. In contrast, DoG fails to accurately find the platelet borders and entirely misses platelets deeper into the thrombus, where the fluorescence intensity decreases. PlateSeg performed well when separating platelets in the low-resolution z axis, where DoG often failed to do so.

**Figure 2.**
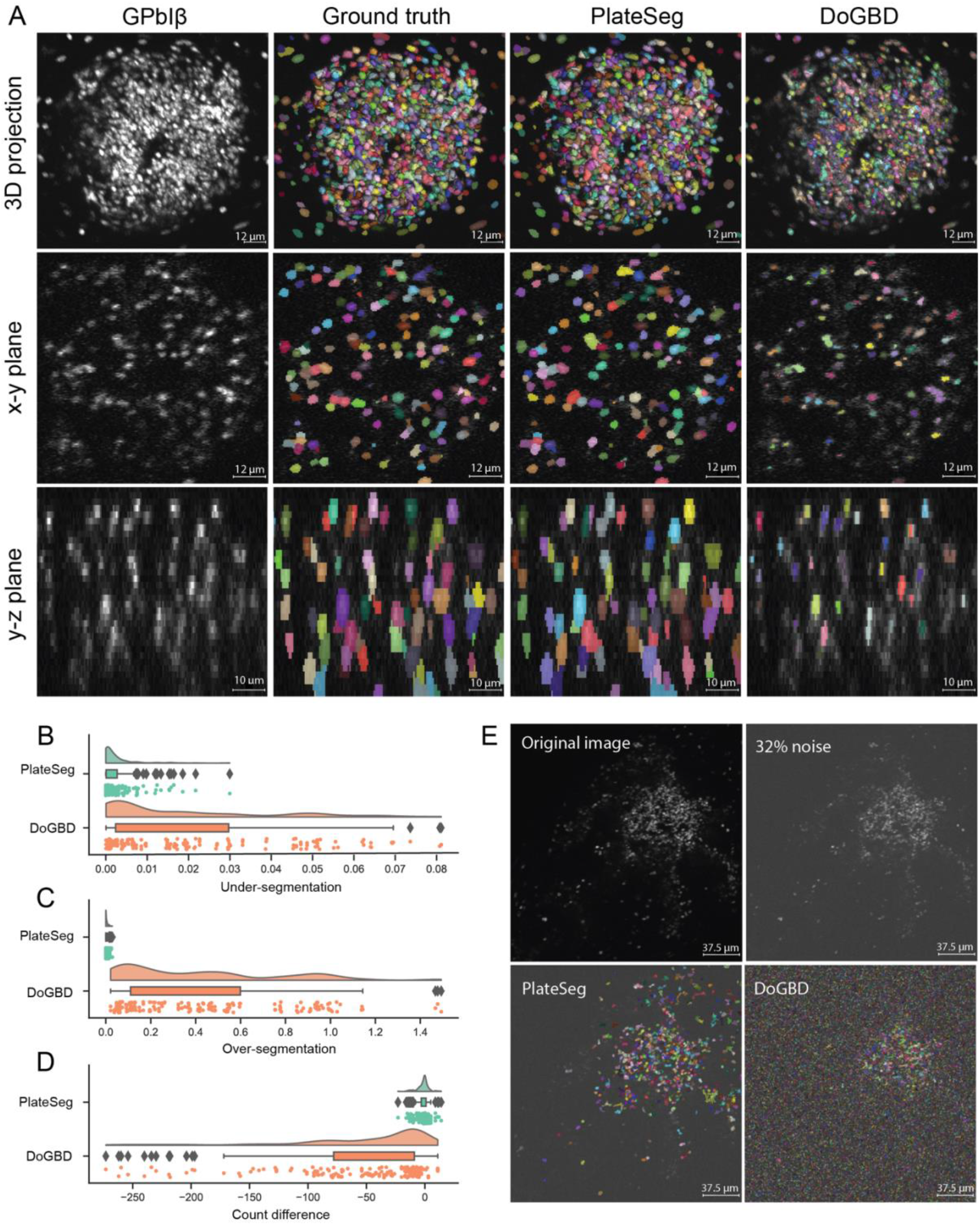
PlateSeg can accurately segment platelets in *in vivo* fractionally stained thrombi. (**A**) Image matrix showing the platelet surface marker GPIbB, ground truth segmentation, PlateSeg segmentation, and DoG segmentation (*from left to right*) in a maximum intensity 3D projection, the xy-plane, and the zy-plane (*from top to bottom*). (**B**) Raincloud plot of under-segmentation in sub-volumes of *in vivo* data processed with PlateSeg and DoG (**C**) Raincloud plot of over-segmentation in sub-volumes of *in vivo* data processed with PlateSeg and DoG. (**D**) Raincloud plot of difference in count between segmentation and ground truth (n_Seg_ – n_GT_) for *in vivo* data processed with PlateSeg and DoG. (**E**) Maximum intensity projection of the original image (*top left*), image with noise of up to 32% of the average image intensity (*top right*), and the noise image overlayed with the PlateSeg segmentation (*bottom left*) and DoG segmentation (*bottom right*).

We used two measures to quantify the accuracy of the automated segmentations by comparing them to the manually-generated labels: variation of information (VI) [23], and a simple count difference. VI has two sub-scores, one measuring the probability that an object will be incorrectly merged with another (under-segmentation) and the other measuring the probability that an object will be incorrectly split (over-segmentation). **Figure 2.B-C** shows these accuracy scores for each sub-volume of image that was input to the network. *PlateSeg* showed very little under-segmentation (0.0025 ± 3.2 × 10^−4^ [mean ± SEM]; 95% CI: 0.0019, 0.0041; **Figure 2.C**), equivalent to only ∼0.3% of objects being incorrectly merged, and little over-segmentation (0.0041 ± 4.4 × 10^−4^; 95% CI: 0.0032, 0.005; **Figure 2.D**), equivalent to only ∼0.4% of objects being incorrectly split. By contrast, the DoG method had higher rates of under-segmentation (0.019 ± 1.5 × 10^−3^; 95% CI: 0.016, 0.022) and very high rates of over-segmentation (0.42 ± 0.026; 95% CI: 0.37, 0.48). This is equivalent to ∼2% of objects being incorrectly merged and ∼42% of objects being incorrectly split (likely due to failure to extend object borders sufficiently). With regards to count difference, *PlateSeg* performed well, missing 1.50 ± 0.36 (95% CI: 0.78, 2.2; **Figure 2.E**) platelets per sub-volume (each sub-volume having 98 ± 5.4 platelets), again substantially outperforming the DoG method, which on average missed 52 ± 4.7 (95% CI: 43, 61).

We also tested PlateSeg and DoG on an image artificially corrupted with noise of up to 32% of image average intensity. PlateSeg showed greater robustness to noise than DoG, which found many objects that did not exist (**Figure 2.D**)

We next applied our segmentation method to image data obtained using an *ex vivo* flow chamber model of thrombosis, performed with either human or mouse blood. We show a representative crop of a segmentation of a thrombus formed from human blood in **Figure 3.A**. *PlateSeg* performs very well on this data, with even lower rates of over-segmentation (6.3 × 10^−4^ ± 1.2 × 10^−5^; 95% CI: 4.0 × 10^−4^, 8.7 × 10^−4^; **Figure 3.B**) and under-segmentation (2.1 × 10^−3^ ± 2.6 × 10^−4^; 95% CI: 1.6 × 10^−3^, 2.7 × 10^−3^; **Figure 3.B**) than seen in *in vivo* data (**Figure 2**). Furthermore, *PlateSeg* made only minimal errors in the count of platelets in a processed crop of image, on average counting 1.3 ± 0.3 cells less the ground truth (95% CI: 0.7, 1.9; **Figure 3.C**), with each sub-volume on average having 76 ± 7 platelets.

**Figure 3.**
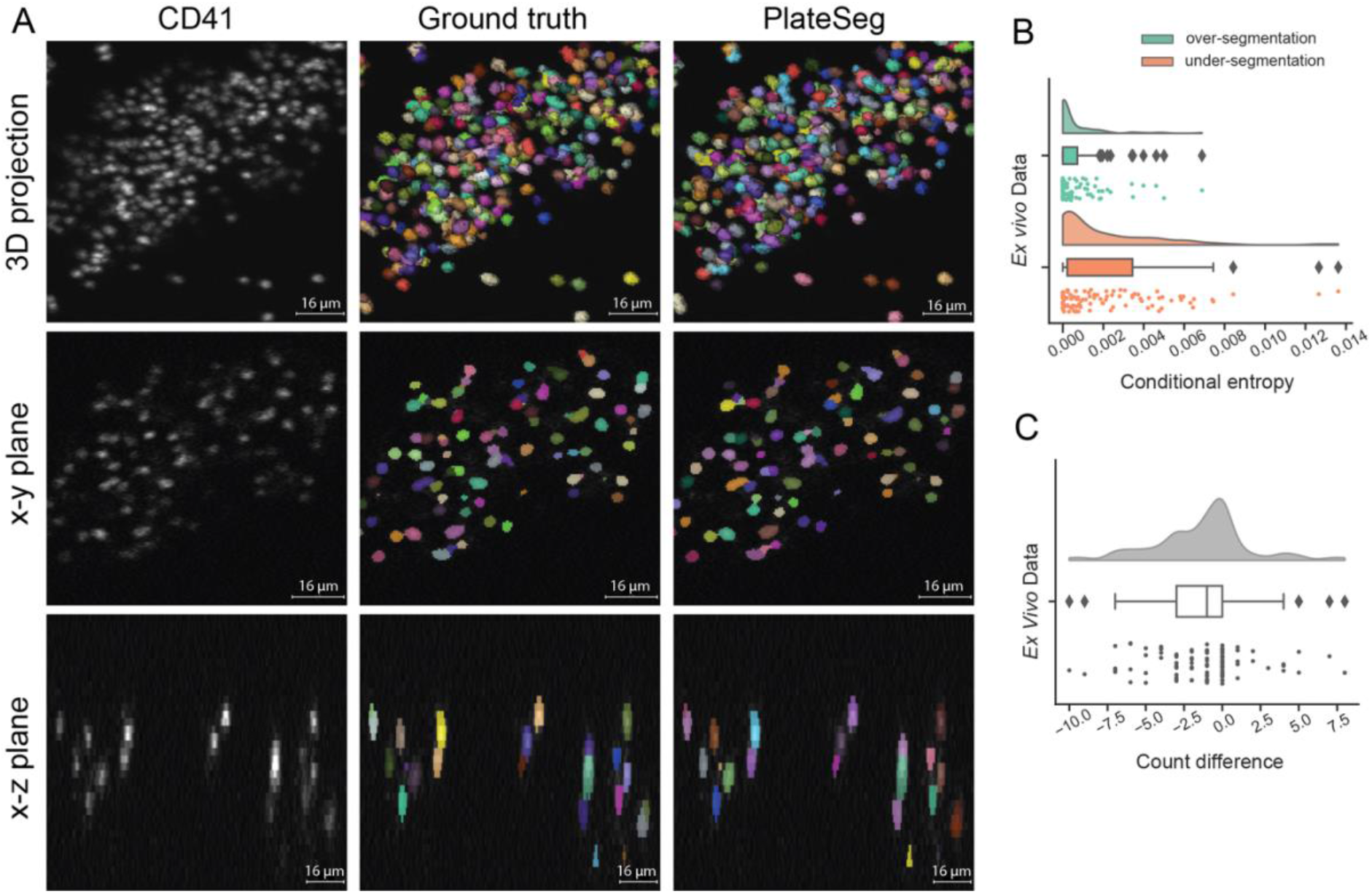
PlateSeg can accurately segment platelets in 3D from confocal z-stacks of time-lapse image data obtained during *ex vivo* thrombus formation. (**A**) Image matrix of platelets fractionally (2.5%) stained with a CD41 surface marker (***left column***) overlaid with ground truth (***middle column***) or PlateSeg (***right column***) segmentations. Platelets can be seen as a maximum intensity projection from below (***top row***), in a slice of the xy-plane (***middle row***) and in a slice of the xz-plane (***bottom row***). This thrombus was formed *ex vivo* from human whole blood flowed over collagen at 600s^-1^. (**B**) Raincloud plots of over-segmentation and under-segmentation in PlateSeg processed image sub-volumes. (**C**) Raincloud plots of count difference between the ground truth and the segmentation (n_Seg_ – n_GT_).

### Platelet tracking in vivo and ex vivo was of high accuracy

Once platelets are segmented, we use a tracking algorithm to follow individual platelets between consecutive timepoints. Representative images showing platelet trajectories are shown in **Figure 4.A** for both *in vivo* (**4.A.1**) and *ex vivo* (**4.A.2**) thrombi. Given the large numbers of platelets tracked (sometimes thousands present in each frame), it is difficult to visualise and assess the quality of a tracking experiment. To facilitate rapid assessment of tracking quality, we developed software to extract a sample of platelet trajectories of up to 30 frames along with a small segment of corresponding image (**Figure 4.B**). Using this software, the user can annotate frames when the trajectory incorrectly jumps to an adjacent platelet (ID swap error) or when a trajectory incorrectly starts or ends (discontinuation errors). We analysed tracking error rates for platelets that become stably incorporated into the thrombus for small (*n =* 64 samples), medium (*n =* 71), and large (*n =* 70) thrombi formed *in vivo* (**Figure 4.C**). Error rates were low for all sizes of thrombi for both ID swap errors and discontinuation errors, occurring in, on average, less than 0.5% of frames. We found that the trajectories from thrombi of different sizes had similar rates of ID swap errors (*H =* 3.1, *p =* 0.21) but different rates of discontinuation errors (*H =* 6.4, *p =* 0.04). This is likely because we did not observe any discontinuation errors in the tracks sampled from the small thrombi.

**Figure 4.**
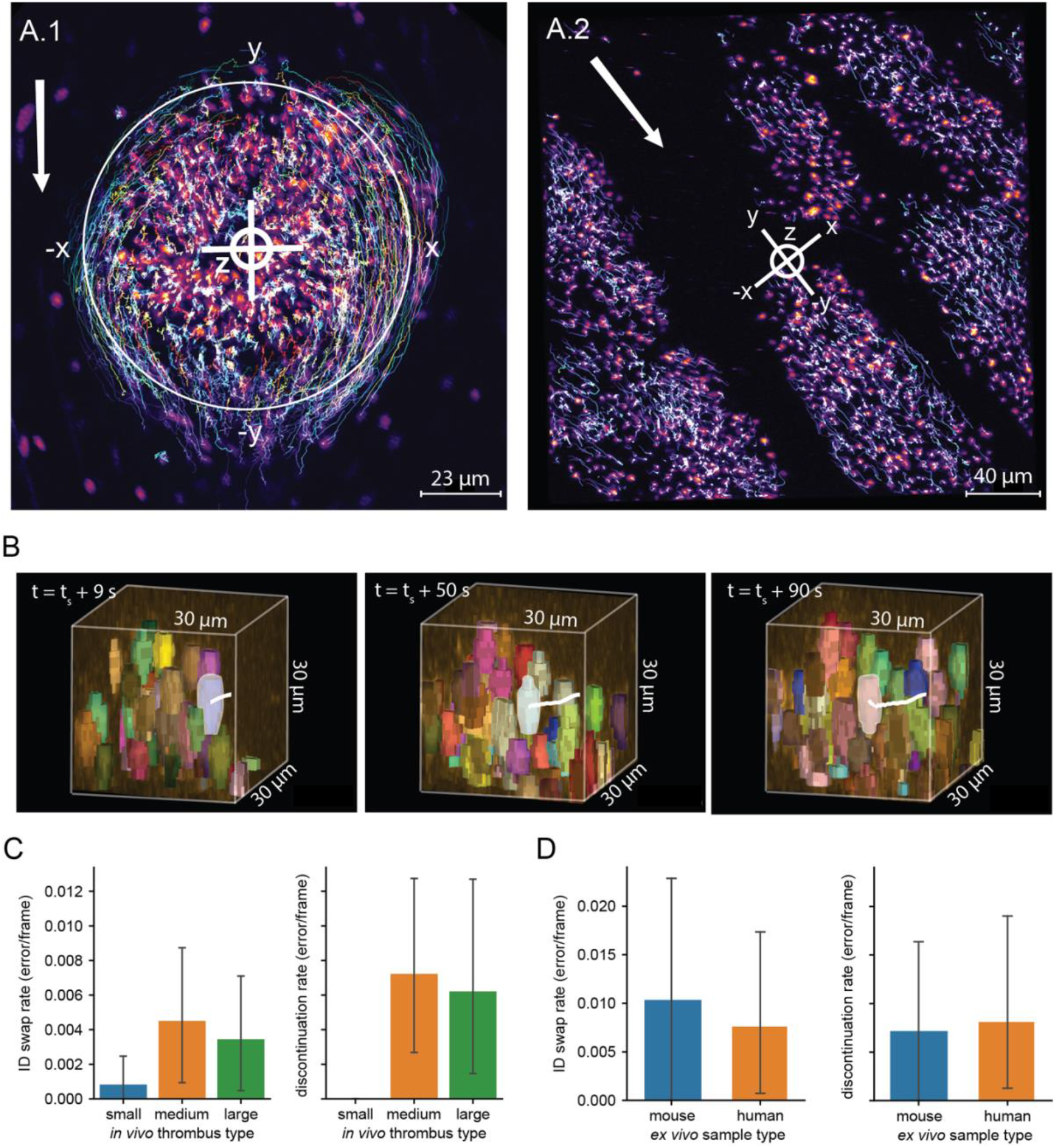
Accurate tracking of platelets in both *in vivo* and *ex vivo* models of thrombus formation using PlateSeg. (**A**) Maximum intensity projection of platelet surface marker GPIbB (displayed using a colour look up table) overlaid with platelet trajectories several minutes after injury in (**A.1**) *in vivo* and (**A.2**) *ex vivo* models of thrombus formation. In these images, ***bright lines*** indicate platelet trajectories over the course of ∼3 min, with warmer colours indicating platelets that stay in the thrombus for longer time periods. The ***crosshairs*** symbol indicates the coordinate axes used to provide platelet coordinates, with letters corresponding to axis labels and polarity. In A.1, the ***large circle*** indicates the region of laser-induced endothelial damage. ***White arrows*** indicate direction of blood flow. (**B**) Visualisation of a single platelet trajectory (highlighted in white) as used for estimating tracking error rates. The platelet is followed for up to 30 frames and frames are annotated where an error occurs. ***Left***: platelet 9 s after the first time point at which it was examined; ***middle***: platelet 50 s after the first time point; ***right***: platelet 90 s after the first time point. (**C**) Tracking error rates in thrombus formation *in vivo* for platelet trajectories longer than 10 frames. ***Left***: rate of ID swap errors, in which the track jumps to an adjacent platelet. ***Right***: rate of discontinuation errors, in which a track incorrectly ends or starts. (**D**) Tracking error rates in thrombus formation *ex vivo* for platelet trajectories longer than 10 frames. ***Left***: rate of ID swap errors, in which the track jumps to an adjacent platelet. ***Right***: rate of discontinuation errors, in which a track incorrectly ends or starts.

We are also able to track platelets in *ex vivo* data with high accuracy. To assess the accuracy of tracking in *ex vivo* tracking analyses we sampled trajectories from experiments with mouse (*n =* 34) and human (*n =* 46) blood. Both ID swaps and discontinuation errors in both mouse and human samples occurred at a rate of around 1% of frames. There was no difference between mouse and human samples for either ID swaps (*U =* 779.0, *p =* 0.96) or discontinuation errors (*U =* 796.5, *p =* 0.79).

### The kinetics of thrombus formation in and ex vivo

Our workflow allows us to quantify aspects of thrombus formation that cannot be measured without segmentation and tracking of platelets. To demonstrate the utility of our method, we describe the kinetics of thrombus formation in our *in vivo* and *ex vivo* models of thrombus formation. **Figure 5.A** shows a 3D rendering of an *in vivo* thrombus (**A.1**), which was formed in mouse mesenteric venules contrasted with *ex vivo* thrombi (**A.2**) from mouse blood formed on collagen in a flow chamber at 600s^-1^. We see that our *in vivo* thrombi accumulate platelets faster (**Figure 5.B**), appear to form dense regions more quickly (**Figure 5.C**), and move more quickly (**Figure 5.D**) than do *ex vivo* thrombi. Nevertheless, under these conditions, the *ex vivo* thrombi gradually attain a similar level of average density to the *in vivo* counterparts.

**Figure 5.**
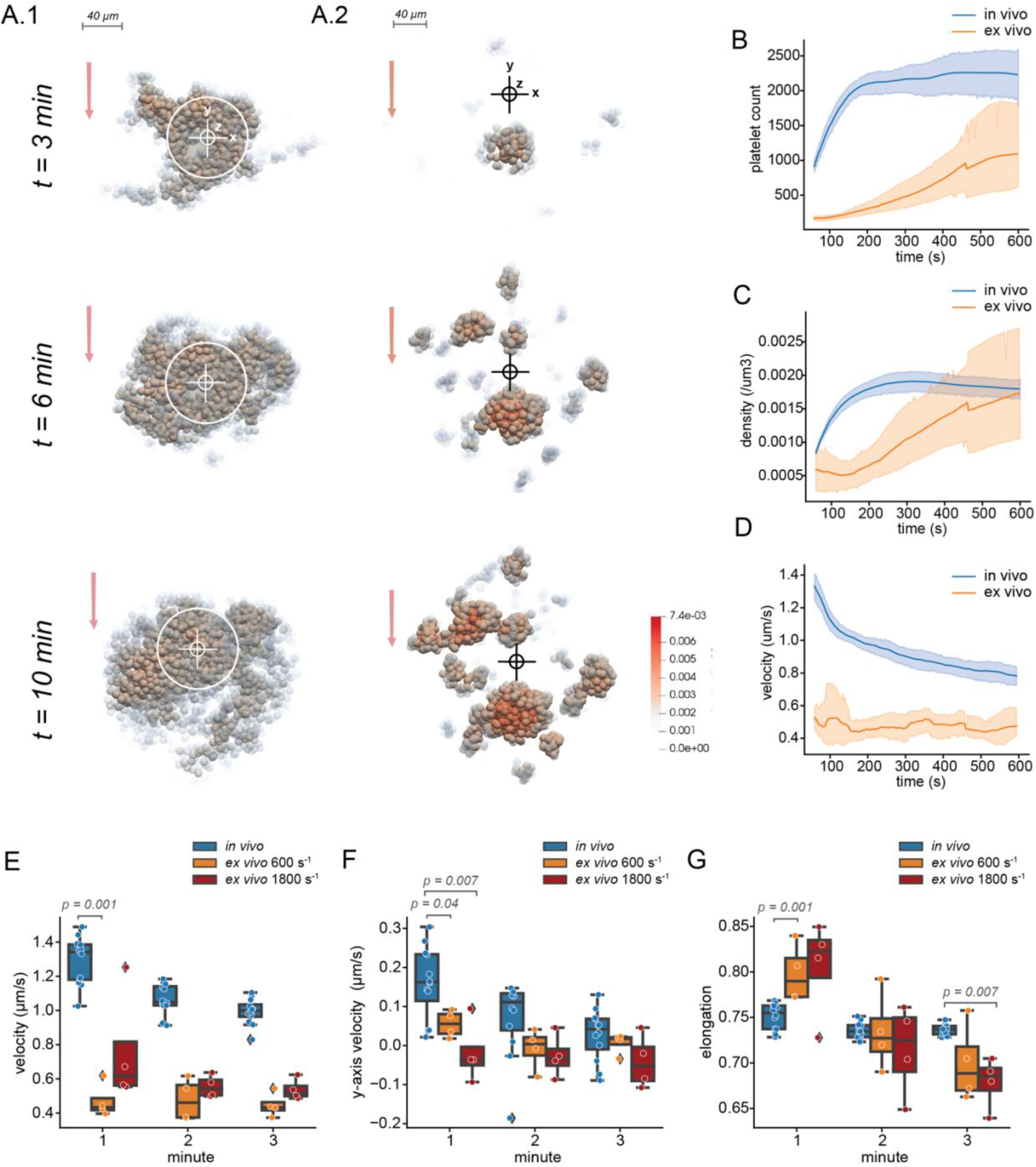
Visualisation and quantification of the kinetics of *in vivo* (in high flow mesenteric venules) and *ex vivo* (flow rate = 600s^-1^) thrombus formation with 2.5% of total platelets stained. (**A**) Rendering of platelet coordinates in (**A.1**) *in vivo* or (**A.2**) *ex vivo* thrombus formation at 3, 6, and 10 min after starting the experiment showing only platelets tracked for more than 10 frames. Each platelet is coloured according to density in the 15μm radius sphere around it. ***White circles*** represent the borders of the laser injury in the *in vivo* thrombus. ***Cross hairs*** show the origin of the coordinate system in each frame. ***Red arrows*** show the direction of blood flow. (**B**) Platelet count in *in vivo* (***blue, n = 12***) and *ex vivo* (***orange, n = 4***) thrombus formation over 10 minutes after starting an experiment. (**C**) Average platelet density in a 15 μm sphere around each platelet in *in vivo* (***blue***) and *ex vivo* (***orange***) thrombus formation over 600 seconds after starting an experiment. (**D**) Average platelet velocity in *in vivo* (***blue***) and *ex vivo* (***orange***) thrombus formation over 600 seconds after starting an experiment. (**E**) Average platelet intra-thrombus velocity in *in vivo* (***blue***) and *ex vivo* (***orange***) thrombus formation at 600s^-1^ and 1800s^-1^ during the first 3 min of the experiment. (**F**) Average platelet intra-thrombus velocity in the axis parallel with blood flow in *in vivo* (***blue***) and *ex vivo* (***orange***) at 600s^-1^ and 1800s^-1^. Here positive values indicate movement against blood flow and negative values indicate passive movement. (**G**) Average platelet elongation in *in vivo* (***blue***) and *ex vivo* (***orange***) at 600s^-1^ and 1800s^-1^. Box plots are annotated with the *p*-values from Mann Whitney U-tests, where these were conducted.

Thrombus formation *in vivo* and *ex vivo* appears to be most different in the first few minutes. We investigated platelet properties in the first three minutes of thrombus formation *in vivo*, and *ex vivo* at 600s^-1^ and 1800s^-1^. In addition to moving more quickly (**Figure 5.E**), *in vivo* thrombi appear to have greater movement against the flow of blood in the first minute for both flow rates (**Figure 5.F**). Interestingly, there appears to be a difference in the average platelet morphology, with platelets in the 600s^-1^ and 1800s^-1^ *ex vivo* thrombi having a more elongated shape at the beginning of experiments (**Figure 5.G**), which may indicate a difference in platelet activation.

## Discussion

In this study we designed an effective and accurate workflow for detecting and tracking large numbers of platelets in both *in vivo* and *ex vivo* models of thrombus formation. Compared to a classical method for object segmentation, our deep learning-based segmentation algorithm performed better segmenting *in vivo* thrombi and was less sensitive to noise [24]. Despite excellent performance, our method remains imperfect, either merging or splitting just under a percent of platelets and missing around 1 in 80 platelets. These errors likely stem from reduced image quality with increasing depth into the blood vessel lumen, which results from optical factors. Here, even expert eyes must carefully inspect the image from multiple angles to correctly label cells. If analyses required higher segmentation accuracy in these regions, this would likely be corrected by further training following time consuming proofreading of low-quality image data. Segmentations of thrombi formed *ex vivo* were more accurate than for those formed *in vivo*. This may be explained by the height of *ex vivo* thrombi, which grow less tall than their *in vivo* counterparts and therefore appear to suffer less from depth-dependent image quality issues.

Using our custom software for error estimation, we showed that platelet tracking using the TrackPy package has low error rates with less than 0.5% of tracks containing ID swaps or discontinuation errors. This software allows users to ascertain the rates of two types of errors: (1) swaps between adjacent platelets and (2) discontinuation errors. Errors have the potential to introduce noise and reduce the validity and power of downstream analysis, but existing tools usually require experts to manually track a complete dataset [25]. This is immensely time consuming and impractical for assessing many different experimental cohorts. By allowing users to subsample and assess platelet tracks, we facilitate error estimation requiring hours rather than days of work. Furthermore, this type of visual analysis may promote a better understanding of the data through hands on interaction, which may help to refine hypotheses and experimental design [10].

Our software computes several measures of platelet behaviour. We used this output to quantify differences between platelet behaviour in *in vivo* and *ex vivo* model of thrombus formation. In *in vivo* model of thrombosis, platelets are exposed to a variety of agonist signals and cues from the damaged blood vessel wall that work synergistically with collagen signalling [26,27]. Perhaps most important, the lack of any anticoagulant in our animal model means that, in contrast to the *ex vivo* system used in this study, the coagulation system is still intact. It is therefore not surprising that our *in vivo* thrombi tended to grow, densify, and move faster than their *ex vivo* counterparts. Platelets in *ex vivo* model of thrombosis were less elongated throughout the experiment, which is consistent with the morphologic analysis under similar flow conditions [28]. Differences in platelet shape between thrombi formed in *ex vivo* and *in vivo* may be explained by the fact that a greater proportion of platelets *ex vivo* become bound to collagen rather than to other platelets. Collagen bound platelets were found to have a more flattened morphology than platelets encountered higher up in the thrombus [29], probably as a consequence of differences in agonist stimulation [30]. Additionally, the hydrodynamic effects of physical thrombus expansion eventually leads to blood flow perturbations [31]. In flow chamber models, larger areas are coated with platelet agonists, which results in the concurrent appearance and growth of multiple thrombi. This may have unpredictable effects on local flow conditions, leading to artefactual variations in platelet activation.

The software we have developed is available through the python package installer and via the napari plugin hub. Our long-term aim for the tracking software is to enable the user to employ other methods for tracking in addition to TrackPy. We also aim to intergrate iterseg with other segmentation methods for better general use. We hope that other researchers will use our software and contribute, either by contributing to our code repository on GitHub or by posting issues and suggestions on the repository.

## Conclusions

We have developed a workflow to accurately find and follow large numbers of individual platelets during thrombus formation. Our methodology can be applied to obtain detailed information from conventional fluorescence microscopy experiments both *in vivo* and *ex vivo* model of thrombosis. The code from our workflow is open source and requires limited Python knowledge to run. Ours is the first report of a deep leaning pipeline for segmentation and tracking of platelets. We aim to continue to maintain and improve the code into the future with the help of other scientists in the community. We hope that our pipeline will provide a versatile and powerful tool for researchers addressing a wide range of questions related to the physiological and pathophysiological roles of platelets.

## Supporting information

Supplementary instructions

