## Supplementary instructions for "Finding and Following: A deep learning-based pipeline for tracking platelets during thrombus formation *in vivo* and *ex vivo*"

### Finding and Following Platelets – A guide to the software

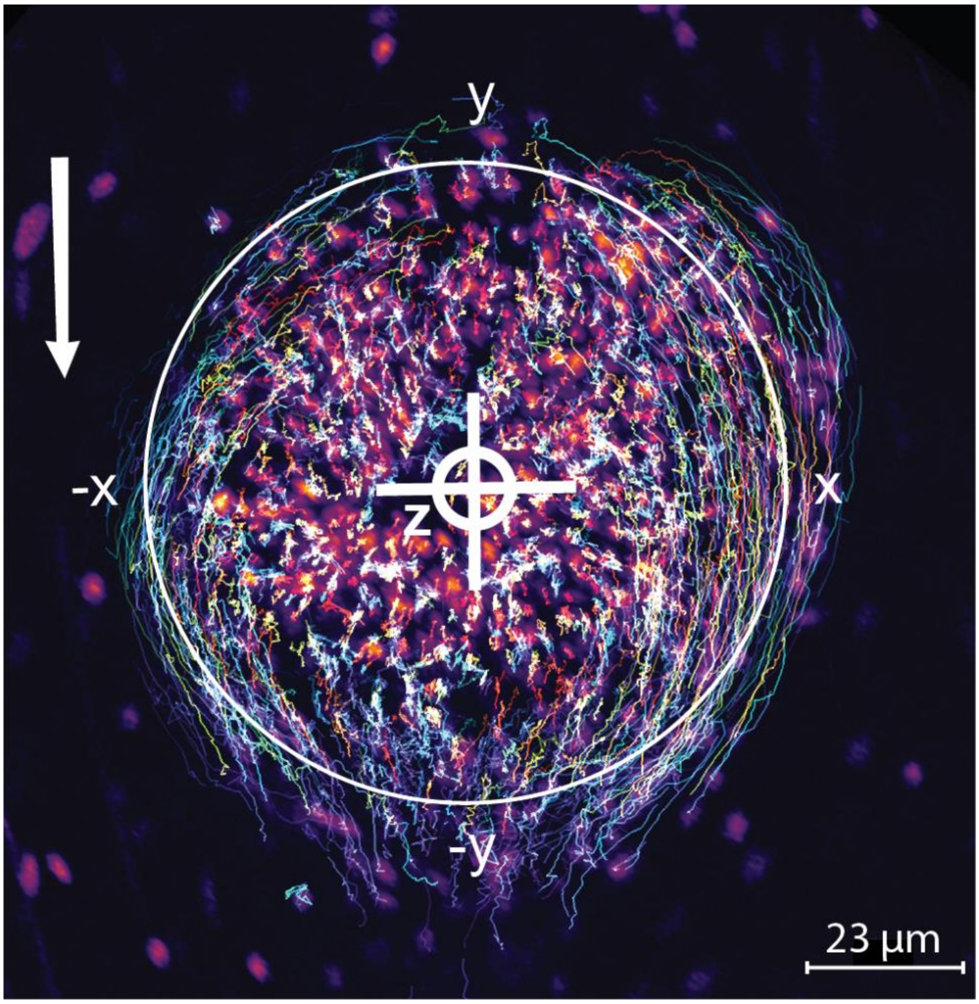


#### *TABLE OF CONTENTS*

**FINDING AND FOLLOWING PLATELETS – A GUIDE TO THE SOFTWARE 1**

**PACKAGE INSTALLATION 2**
**DOWNLOADING THE SAMPLE DATA 4**
**SEGMENTING SINGLE IMAGES USING ITERSEG 6**
**SEGMENTATION AND TRACKING WITH ITERSEG AND PLATETRACK 7**
**ANNOTATING TRACKS WITH ANNOTRACK 9**
**TRAINING A U-NET FOR AFFINITIES-BASED SEGMENTATION 12**

#### Package installation

1. Make sure your computer has a Python environment manager such as *Anaconda* (https://www.anaconda.com/download) or *miniforge* ([https://conda-forge.org/miniforge/](https://conda-forge.org/miniforge/)) installed.
2. Open terminal (if you are using MacOS or Ubuntu) or anaconda prompt or similar command line (if you are using Windows).
3. To create a new environment, type the following: `conda create --name p10 python=3.10 pip` and press enter. We want to create a new environment because some of the packages we depend on (e.g., PyTorch) don't always play well in the sandpit with other packages that could be installed in an existing environment. This sort of interaction would cause errors when running the program. To activate the environment type: `conda activate p10` then press enter.

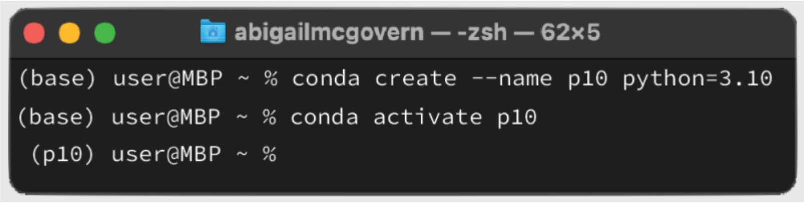


4. To install the plugins, type: `pip install "napari[pyqt5]" iterseg platetrack annotrack` and press enter.

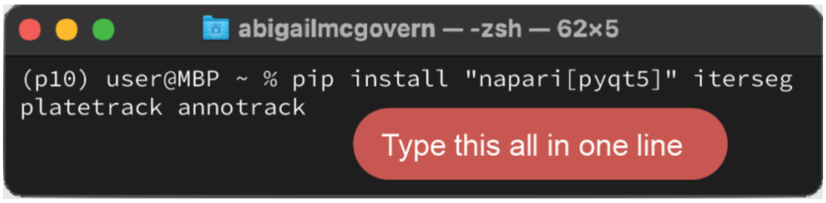


5. To open napari, type: `napari` and press enter. The graphical user interface will look as shown below. You can access the plugins through the plugins menu. When you hover over each plugin, you can see any functions it offers.

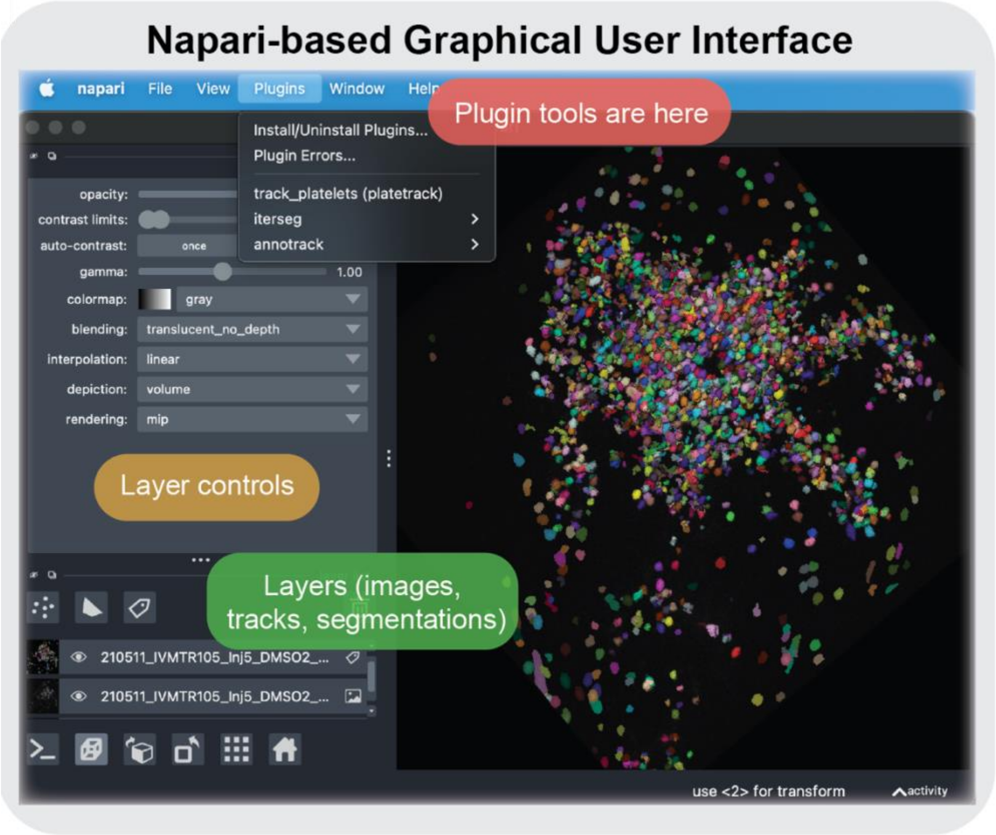
Napari-based Graphical User Interface
napari File View Plugins Window Help
Plugin tools are here
Install/Uninstall Plugins...
Plugin Errors...
track_platelets (platetrack)
iterseg
annotrack
opacity:
contrast limits:
auto-contrast:
once
gamma:
1.00
colormap:
gray
blending:
translucent_no_depth
interpolation:
linear
depiction:
volume
rendering:
mip
Layer controls
Layers (images, tracks, segmentations)
210511_IVMTR105_Inj5_DMSO2_...
210511_IVMTR105_Inj5_DMSO2_...
use <2> for transform
activity

#### Downloading the sample data

Our sample data can be downloaded at the following link:

https://www.dropbox.com/scl/fo/zplk5nxyaqmjjaisldvr6/h?rlkey=cy7rzy3e0pct4gojypsojmvyv&dl=0 [**Note to editor**: to be replaced with figshare DOI]

A description of the folder structure and contents is show below (folders are in **BOLD**; note that the description goes over the page):

- **Segmentation**
  - **exvivo_example**
    - **ground_truth**: 3D ground truth segmentations for individual frames of *ex vivo* data. Data are of dimensions (x, y, z = 512, 512, 33). The last two letters in the file name correspond to the initials of the proof-reader. The number in the file name corresponds to the number of the
    - **images**: 3D images corresponding to the ex vivo ground truth data for individual image frames. Images show the platelet surface marker. Data are of dimensions (x, y, z = 512, 512, 33).

  - **invivo_example**
    - **ground_truth**
      - **large – DMSO**: 3D ground truth segmentations for individual frames of data from a large DMSO treated thrombus *in vivo*. Data are of dimensions (x, y, z = 512, 512, 33).
      - **medium – control**: 3D ground truth segmentations for individual frames of data from a medium size control thrombus *in vivo*. Data are of dimensions (x, y, z = 512, 512, 33).
      - **small – cangrelor**: ground truth segmentations for individual frames of data from a small cangrelor treated thrombus *in vivo*. Data are of dimensions (x, y, z = 512, 512, 33).

        - **images**: images for individual frames of data from the small, medium, and large thrombi described above. Images show the platelet surface marker. Data are of dimensions (x, y, z = 512, 512, 33).

    - **training_data**
        - **training_gt**: A series of ground truth segmentations which can be used for training a network. In this folder, there are thee labels from an early in an *ex vivo* flow experiment and seven images from *in vivo* experiments.
        - **training_images**: A series images which can be used, in conjunction with the ground truth segmentations above, for training a network. The images show platelet surface marker.

- **Tracking**
    - **human_exvivo**: contains two 10-minute time series featuring thrombi formed from human whole blood growing under different flow conditions (3000 $s^{-1}$ and 600 $s^{-1}$). These images have four channels: (1) a calcium marker, (2) a fibrin marker, (3) a platelet surface marker, and (4) a TD image. The platelet surface marker channel is the one that should be used for segmentation.
    - **mouse_exvivo**: contains two 10-minute time series featuring thrombi formed from mouse whole blood growing under different flow conditions (1800 $s^{-1}$ and 600 $s^{-1}$). These images have four channels: (1) a calcium marker, (2) a fibrin marker, (3) a platelet surface marker, and (4) a TD image. The platelet surface marker channel is the one that should be used for segmentation.
    - **mouse_invivo**: contains two 10-minute time series featuring thrombi formed from mouse whole blood growing under different flow conditions (1800 $s^{-1}$ and 600 $s^{-1}$). These images have four channels: (1) a calcium marker, (2) a p-selectin marker, (3) a platelet surface marker, and (4) a TD image. The platelet surface marker channel is the one that should be used for segmentation.

#### Segmenting single images using Iterseg

1. First hover over the iterseg plugin in the plugins menu. You will see a widget called ***load_data***. Click on this to bring up the widget to load the data. We want to load in an image or group of images to segment (in either the **segmentation > exvivo_example > images** or **segmentation > invivo_example > images** folder). Choose a name for your image layer in the layer name box. You can load a single zarr image by clicking the ***choose directory*** button and selecting the zarr file. If you wish to load all of the images in a folder select that folder (e.g., or **segmentation > invivo_example > images**).  Set the scale to (2, 0.5, 0.5). Make sure the layer type is Image and the data type is individual frames. Make sure split channels is not ticked. Click ***Run***. Once the image is loaded go to autocontrast and click once to adjust the contrast and click the cube ($2^{nd}$ icon in) in the bottom left corner to make the image 3D.

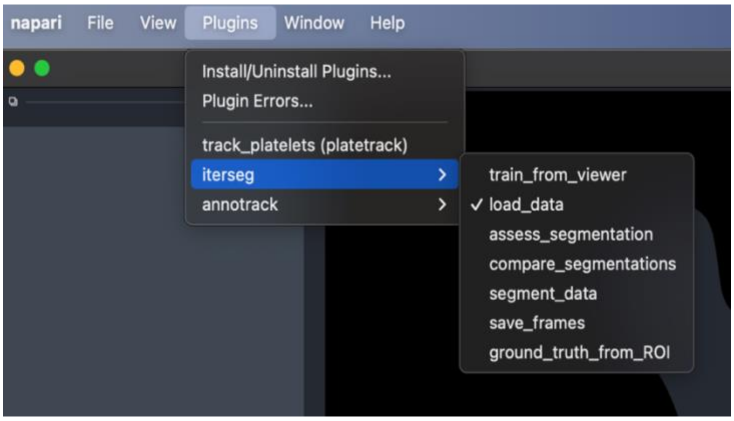

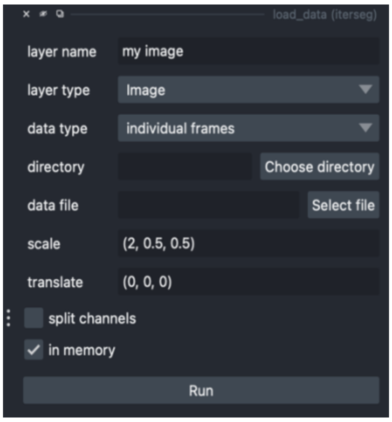


2. Go back to the plugins menu and click ***segment_data***. Make sure the input volume layer is set to the image you just loaded. Choose a directory that you want to save the segmentation into by clicking ***choose directory*** and choose a name for the save file (you do not need to include an extension .zarr will be added). Make sure the segmenter is set to ***affinity-unet-watershed***. You do not need to choose a network or configuration file, for now we will use the default version. As we train the network on extra data, we will make other unets available. Layer reference should be None, chunk size should be (10, 256, 256) and margin should be (1, 64, 64). Make sure debug is unticked. To segment the image or images, click ***Run***.

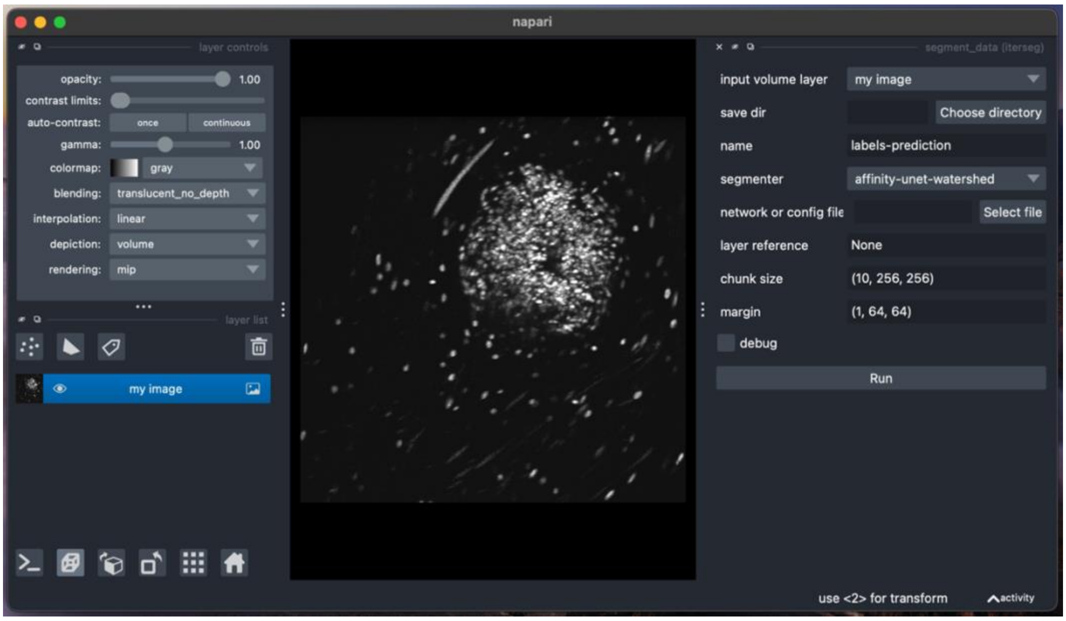


#### Segmentation and tracking with Iterseg and Platetrack

1. To segment a full timeseries (video of a thrombus) with Iterseg, follow the same process as above but choose one of the full timeseries from one of the tracking folders using the ***Choose directory*** button (**tracking > human_exvivo**, **tracking > mouse_exvivo**, or **tracking > mouse_invivo**). The only difference in loading this data is that this time you need to make sure <u>split channels is ticked</u>. For these images channel 0 contains a calcium marker, channel 1 ( <*layer name*> [1])contains a fibrin marker, and channel 2 (<*layer name*> [2]) contains the surface marker that you will use for the segmentation. Repeat step 2 from above ensuring to choose the channel with the surface marker. This process may take several minutes to several hours depending on your computer.

2. Once the segmentation is complete open the ***track_platelets (platetrack)*** widget. Once the widget loads, choose the segmentation layer as the labels layer. Tick use all image layers (the image layer chosen doesn't matter once this is ticked). Type a sample name and a treatment name. Leave the microns as they are set. Choose a directory to save the output to by clicking ***Choose directory***. Leave the save file blank (you would use this to add to an existing file). Change the save format to csv (unless you prefer parquet). Leave the other settings as they are and Click ***Run***.

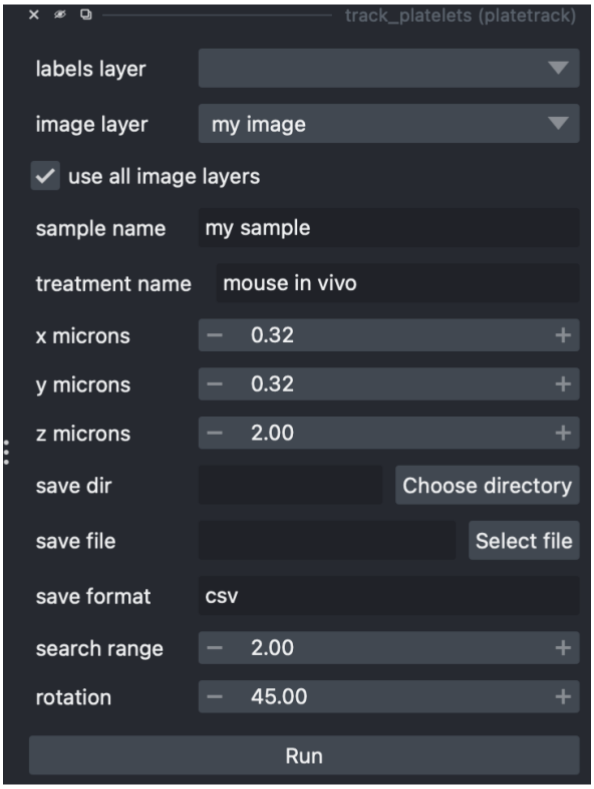

#### Annotating tracks with Annotrack

1. To generate track samples and annotate them, you will need to first generate a csv file (comma separated variable file) with the paths to the images, tracks, and labels in the columns. A good way to make this csv is by using excel to make a spreadsheet (example below) and save as a csv. The example below shows the required columns for the csv. **We you can find a template csv at the link provided ([https://gist.github.com/AbigailMcGovern/975ed4dd686971c99ca6970f12e6f0bb](https://gist.github.com/AbigailMcGovern/975ed4dd686971c99ca6970f12e6f0bb)), which you can download and fill in on excel (or pages)**. The first three columns are called *image_path*, *labels_path*, *tracks_path*, and in these you will add the file paths. A file path tells the computer which folder to look in for the file. Try to avoid any special characters when naming the folders and files – this will help you to avoid errors. You get a file path on a Mac by right clicking the file then clicking on Get Info. Copy and paste the entry next to "*Where*:" then copy the entry under "*Name & Extension*:". The next columns *n_samples* and *sample_type* allow you to specify the number of track samples to take from each file and the category (e.g., treatment, specific antibody, shear rate, laser power, you name it), respectively. NB: The n_samples column is optional (see step 2).

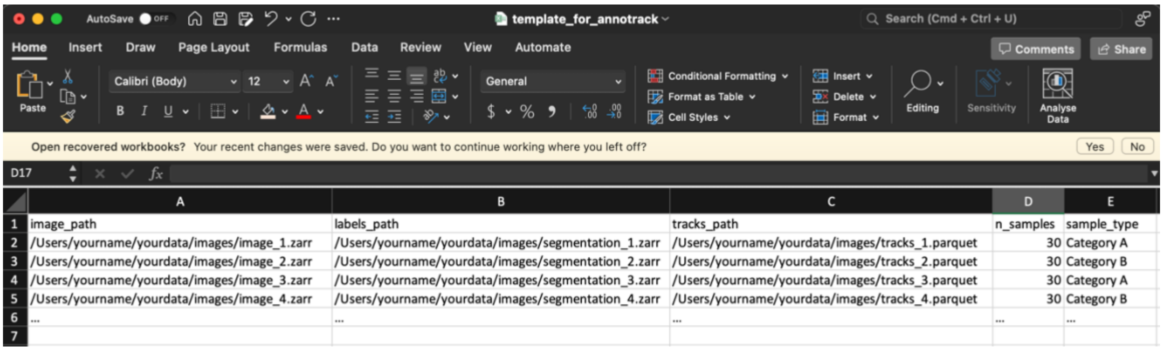

| | A | B | C | D | E |
|---|---|---|---|---|---|
| 1 | image_path | labels_path | tracks_path | n_samples | sample_type |
| 2 | /Users/yourname/yourdata/images/image_1.zarr | /Users/yourname/yourdata/images/segmentation_1.zarr | /Users/yourname/yourdata/images/tracks_1.parquet | 30 | Category A |
| 3 | /Users/yourname/yourdata/images/image_2.zarr | /Users/yourname/yourdata/images/segmentation_2.zarr | /Users/yourname/yourdata/images/tracks_2.parquet | 30 | Category B |
| 4 | /Users/yourname/yourdata/images/image_3.zarr | /Users/yourname/yourdata/images/segmentation_3.zarr | /Users/yourname/yourdata/images/tracks_3.parquet | 30 | Category A |
| 5 | /Users/yourname/yourdata/images/image_4.zarr | /Users/yourname/yourdata/images/segmentation_4.zarr | /Users/yourname/yourdata/images/tracks_4.parquet | 30 | Category B |
| 6 | ... | ... | ... | ... | ... |
| 7 | | | | | |

2. Once you have created the csv, you only need to load the annotrack widget ***sample_from_csv*** and fill in the fields (as shown below). In the first field, click ***Select file*** to choose the csv file you just created. In the next click ***Choose directory*** to choose the folder into which to save the file. Now you can give the output file a name (this is the file into which all of your annotations will be saved). Leave the

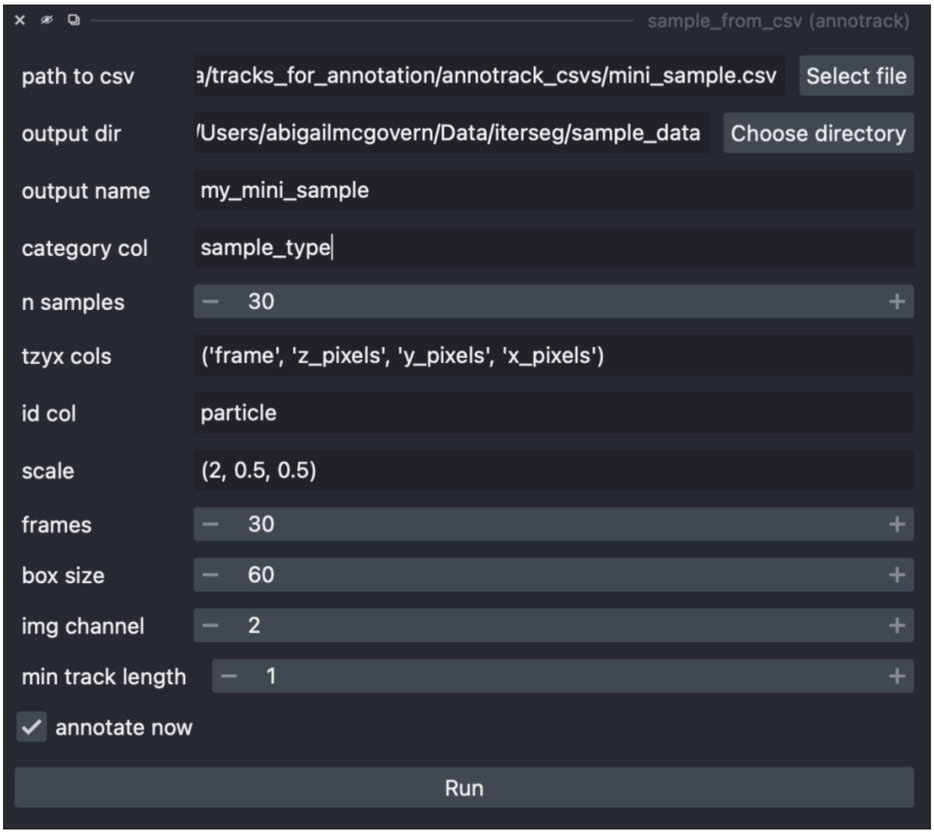


category col as *sample_col* (unless you changed it in the csv, in which case they need to match). Leave the *n samples* box as *30* unless you have (1) not included the column in your csv and (2) want to take a different number of samples from each tracked experiment. If you have included the n_samples column in your csv this will override the *n samples* box here. Leave *tzyx cols* as *('frame', 'z_pixels', 'y_pixels', 'x_pixels')* and *leave id* col as *particle* unless you have changed column names in your tracks files. These values correspond to the names of the columns containing information about frame, coordinates in pixels, and platelet ID (and the order is important). For our data set scale should be (2, 0.5, 0.5) as this is the size in microns of the pixels in z, y, and x coordinate axes. Leave the *frames*, *box size*, and *img channel* fields as *30*, *60*, and *2*, respectively for optimal viewing. The *frames* parameter indicates the maximum number of frames you will be shown for each track segment. The *box size* parameter indicates the size of the bounding box you will be shown around the tracked platelet in microns. *Img channel* is the channel with the platelet surface marker (which you will use to help annotate errors). *Min track length* is the minimum track length of platelets you will be shown. *Min track length* of 1 indicates you will look at all tracked platelets. However, if you are interested in looking at longer tracks (i.e., stably incorporated platelets), select a higher number.

3. Click Run to annotate the tracks. To annotate the tracks use the following key strokes:

   a. Keys to navigate and annotate samples
   b. '2' - move to next sample
   c. '1' - move to previous sample
   d. 'y' - annotate as correct (will move to the next sample automatically)
   e. 'n' - annotate as containing an error (will move to the next sample automatically)
   f. 'i' - annotate the frame following a ID swap error
   g. 't' - annotate the fame following an incorrect termination
   h. 'Shift-t' - annotate the frame containing a false start error
   i. 's' - annotate an error ('i', 't', or 'Shift-t') as being associated with a segmentation error (merge or split of objects)

   When an error is associated the specific frame ('i', 't', 'Shift-t', or 's'), the frame number (within the original image) will be added to a list of errors for the sample within the sample's (.smpl) info data frame. E.g., you may have a list of ID swaps for your sampled track segment ([108, 111, 112]) and a corresponding list of segmentation error associations ([108, 112]).

#### Training a U-net for affinities-based segmentation

1. Before trying to train a new neural network on your own data, we recommend you run through training one on our sample data. The data with which to train the network can be found in the ***segmentation > training_data*** folder. Use the ***load_data*** widget (as you would when segmenting data) to load in the stack of training images and ground truth segmentations. Load the images by selecting layer type of *Image* and then click Choose directory to choose the ***segmentation > training_data > training_images*** folder. After giving the layer a name click ***Run***. Then load the ground truth by selecting layer type of *Labels* and then click Choose directory to choose the ***segmentation > training_data > training_gt*** folder. After giving the layer a name again click ***Run***.

2. Once you have loaded in the data you will be able to train the network using the ***train_from_viewer*** widget (shown below). Once you have loaded the widget first make sure the correct image and labels layers are selected. Then choose a directory into which to save output using ***Choose directory*** (includes: checkpoints, the unet, training chucks, computed training labels, etc). Set the ***scale*** to *(2, 0.5, 0.5)*. The ***mask prediction*** allows you to choose what type of feature channel is fed to the segmentation algorithm to find platelet foreground pixels – *mask* is the best option. The ***centre prediction*** function allows you to choose what type of feature channel is fed to the segmentation algorithm to find the centre of platelets. For this I would suggest *centreness*, although *centreness-log* produces similar results. Set ***affinities extent*** to *1*. Give your output U-net a name in ***training name***. The remaining parameters determine the training program used. Use *BCELoss* as the loss function for this data. Default parameters are ***learning rate*** *= 0.01*, ***epochs*** *= 4*, ***validation split*** *= 0.2* (20%), and ***n each*** *= 50*. Learning rate determines how much the values in the network (which determine its output) are updated (corrected with respect to the ground truth) with every step of training. Epochs refers to the number of times you cycle through the data. Validation split determines how much of the data is used for validation versus learning (usually 20%). N each is how many subsamples are taken and augmented from the training data (n each x n samples = mini batch size – in this

case 450). If you have ***predict labels*** ticked, then the trained network will be used to produce a segmentation of the training data once training is done (this will be displayed as a new layer). If you have ***save labels*** ticked the newly predicted labels will be saved. Whist predicting on training data is not a particularly valid way to assess segmentation quality, it will give you a good indication of how the network is performing.

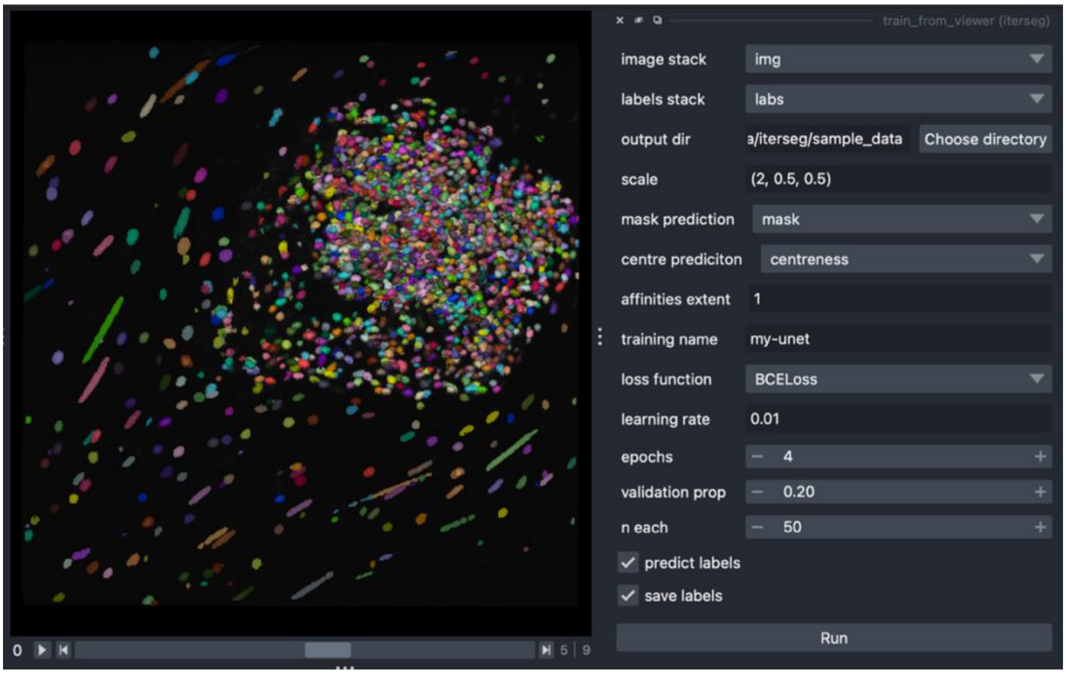


3. Once you are happy with the parameters, click ***Run***. Once running, the results of training will be printed in the command line or terminal you used to launch napari. The command line will show you (1) the progress though training (might take several hours depending on your computer), and (2) the training and validation loss throughout training.

4. In the output in the directory you choose, you will be able to open a text file called unet_paths.txt. This is a plain text file that will record the file paths to the u-net from any training session for which that output directory was chosen. This will tell you where the unet is. The file path for the unet can be pasted into (or found using the ***Choose file*** button) the ***unet or config file*** field of the ***segment data*** widget to use the unet to segment with.
